# Antibacterial effectors in *Dictyostelium discoideum*: specific activity against different bacterial species

**DOI:** 10.1101/2024.05.30.596688

**Authors:** Raphael Munoz-Ruiz, Otmane Lamrabet, Tania Jauslin, Cyril Guilhen, Alixia Bourbon, Pierre Cosson

## Abstract

*Dictyostelium discoideum* is a phagocytic amoeba continuously eating, killing and digesting bacteria. Previous studies have detected in *D. discoideum* cell extracts a bacteriolytic activity effective against *Klebsiella pneumoniae* bacteria.

In this study we characterized bacteriolytic activities found in *D. discoideum* cell extracts against five different bacteria (*K. pneumoniae*, *Escherichia coli*, *Pseudomonas aeruginosa*, *Staphylococcus aureus*, and *Bacillus subtilis).* We first analyzed the bacteriolytic activity against these five bacteria in parallel over a range of pH values. We then measured the remaining bacteriolytic activity in *D. discoideum kil1* and *modA* KO mutants. We also performed partial fractionation of *D. discoideum* extracts and assessed activity against different bacteria. Together our results indicate that optimal bacteriolytic activity against different bacteria results from the action of different effectors. Proteomic analysis allowed us to propose a list of potential bacteriolytic effectors.

**IMPORTANCE:** Many antibacterial effectors have been characterized over the last decades, and their biological importance, mode of action and specificity is often still under study. Here we characterized *in vitro* bacteriolytic activity in *D. discoideum* extracts against five species of gram-negative and gram-positive bacteria. Our results reveal that optimal lysis of different bacteria mobilizes different effectors. Proteomic analysis generated a list of potential bacteriolytic effectors. This work opens the way for future analysis of the role of individual effectors in living *D. discoideum* cells.

## INTRODUCTION

Phagocytic cells ingest bacteria, then kill and destroy them in phagosomes. Intracellular killing and destruction of bacteria in phagosomes is a complex multistep process. This process has been extensively studied in mammalian professional phagocytic cells (neutrophils and macrophages), which defend the body against bacterial infections. A large collection of molecular mechanisms has been proposed to play a role in intracellular killing and can be grouped in four general categories [1]: toxic reactive oxygen or nitrogen species, specific ions (including protons), peptides capable of permeabilizing bacterial membranes, and enzymes digesting the bacterial constituents.

*Dictyostelium discoideum* amoebae are found in forest soil, where they feed upon other soil microorganisms, in particular bacteria [2]. *D. discoideum* cells are easy to handle and their haploid genome is relatively easy to manipulate. *D. discoideum* has been used as a model to study many unicellular traits (e.g. motility, chemotaxis or phagocytosis) as well as its starvation-induced multicellular development [3]. More specifically, *D. discoideum* is a model phagocytic cell to study how phagocytic cells ingest, kill and destroy bacteria in phagosomes. We analyzed previously the phenotypes of a collection of *D. discoideum* mutants and showed that different *D. discoideum* gene products are required to destroy different bacterial species in living cells [4]. Our results also demonstrated that bacterial death in phagosomes precedes the permeabilization of its membrane(s) and its gradual dismantling by digestive enzymes [5].

Both in mammalian phagocytes and in *D. discoideum*, the list of known phagosomal bactericidal mechanisms is presumably incomplete, as well as their relative importance at different stages of the bacteriolytic process. It is also unclear whether different mechanisms are at play to ensure killing of different bacteria.

Antibacterial mechanisms can be studied following at least two distinct approaches. First, cellular gene products potentially implicated in intracellular killing can be genetically altered to evaluate their importance *in vivo*. Second, proteins with antibacterial activity can be identified *in vitro* in cellular extracts, purified and characterized.

In *D. discoideum*, we showed previously that *D. discoideum* cell extracts exhibit a bacteriolytic activity against *K. pneumoniae* bacteria [6]. A partially purified *D. discoideum* bacteriolytic fraction contained at least 16 proteins. In the current study, we analyzed the bacteriolytic activity of *D. discoideum* extracts against several gram-negative and gram-positive bacteria. Besides providing an extensive analysis of potential antibacterial proteins, our results indicate that optimal lysis of each of the five bacteria mobilizes different cellular proteins.

## RESULTS

### *D. discoideum* extracts exhibit bacteriolytic activity against five different bacteria

We first assessed in parallel the bacteriolytic effect of *D. discoideum* extracts against three gram-negative bacteria (*K. pneumoniae*, *E. coli* and *P. aeruginosa*), as well as two gram-positive bacteria (*S. aureus* and *B. subtilis*). In order to compare the ability of *D. discoideum* cell extracts to lyse different types of bacteria, we mixed *D. discoideum* cell extracts with comparable numbers of bacteria of each bacterial species and visualized the bacteria after 2 hours. Some bacteria (*K. pneumoniae* and *P. aeruginosa*) are easily visualized and counted, and they were both lysed with comparable efficiencies (Fig. 1A). For other bacteria (*E. coli*, *B. subtilis* and to a much larger extent *S. aureus*), the presence of large bacterial aggregates prevented a precise counting of bacteria (Fig. 1A). However, in all cases it was apparent that exposure to *D. discoideum* extracts resulted in a significant cell lysis. The amount of lysis was quantified by counting in each picture the surface occupied by unlysed bacteria (Fig. 1B). This quantification confirmed that *D. discoideum* cell extracts caused massive lysis of all five bacteria used in this study. Bacteria that aggregated were apparently less efficiently lysed than non-aggregating bacteria. This could reflect the fact that bacteria included in aggregates were less accessible to bacteriolytic effectors. Alternatively, the quantification method may underestimate the efficiency of lysis when applied to bacterial aggregates. Overall, it appears that different bacteria were lysed with comparable efficiencies by *D. discoideum* cell extracts. From a practical point of view, assessing the optical density of the bacterial suspension provided a measure of bacterial lysis consistent with microscopical analysis (Fig. 1C). Indeed, the optical density of the bacterial suspension decreased gradually as were lysed. This method was used in the rest of this study to determine the bacteriolytic activity of different cellular extracts.

**Figure 1.**
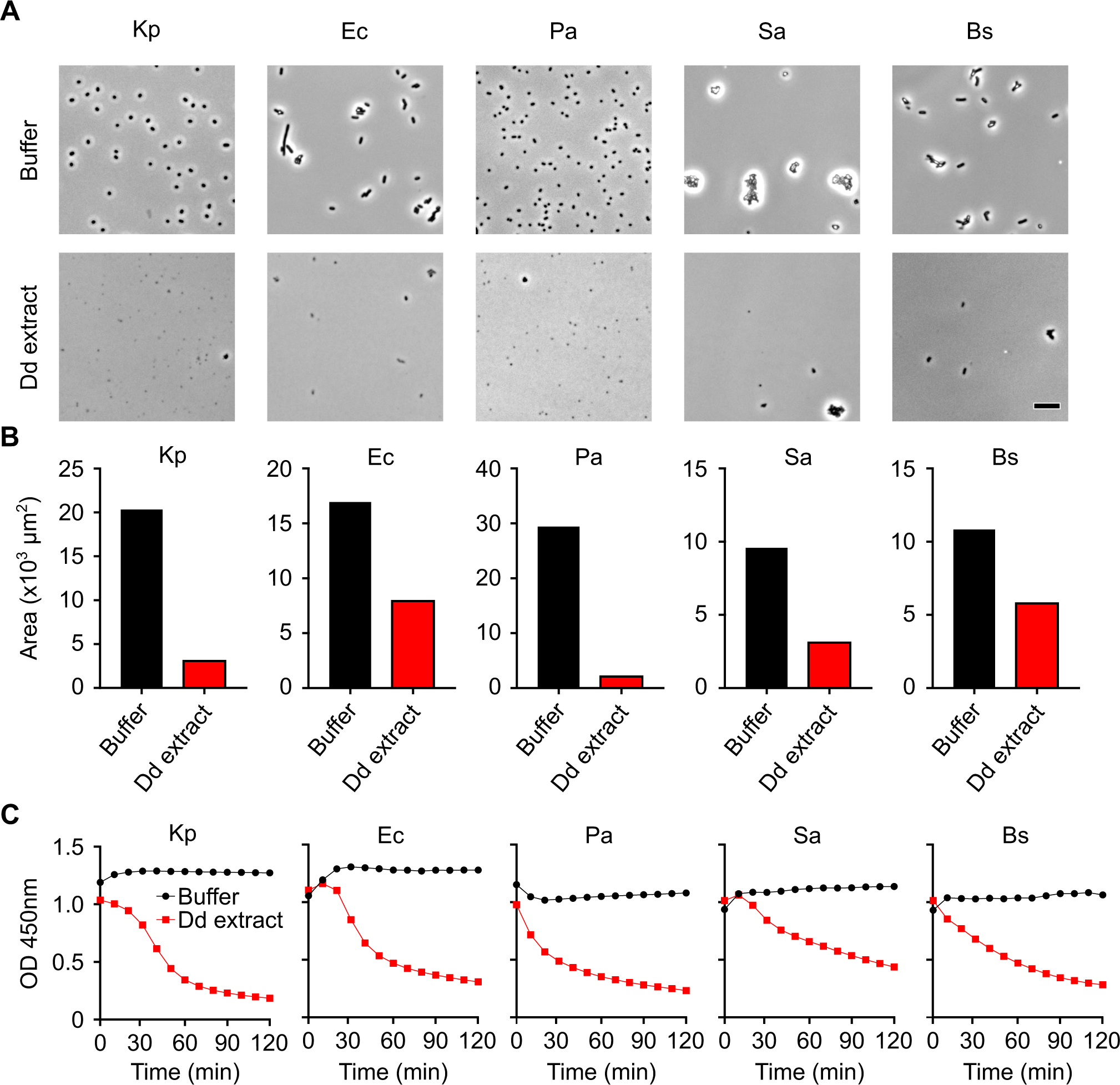
Assessing the bacteriolytic activity of D. discoideum extracts against different species of bacteria. **A**. Different bacteria were incubated for two hours in the presence (Dd extract) or absence (Buffer) of unfractionated D. discoideum extracts. Bacterial lysis was assessed visually using phase contrast microscopy. Kp: K. pneumoniae; Ec: E. coli; Pa: P. aeruginosa; Sa: S. aureus ; Bs : B. subtilis. **B**. The area occupied by bacteria in pictures was quantified to evaluate the degree of bacterial lysis. **C**. Bacterial lysis was followed by measuring the optical absorbance (OD) at 450 nm. Extracts from D. discoideum lysed all bacteria tested with similar efficiencies, and the lysis was most easily followed by measuring optical density. Dd: D. discoideum.

To obtain more quantitative estimates of bacteriolytic activity, bacterial lysis was quantified by testing the activity of several dilutions of *D. discoideum* extracts. In the example shown, the effect of pure or diluted *D. discoideum* extracts was tested on *E. coli* at pH 2 (Fig. 2A). As expected, bacterial lysis was less efficient as the cell extract was diluted and no lysis was observed in buffer alone. We first used this assay to assess bacteriolytic activities over a range of pH. As previously described [6], *D. discoideum* extracts lysed efficiently *K. pneumoniae* at pH 1-2 (Fig. 2B). This very acidic pH reproduces the conditions found in the very acidic *D. discoideum* lysosomes and phago-lysosomes [7, 8]. The bacteriolytic activity against *E. coli* (Fig. 2C), *P. aeruginosa (*Fig. 2D), *S. aureus* (Fig. 2E) and *B. subtilis* (Fig. 2F) was also optimal at very acidic pH (1.5-2). However, the activity profiles differed when different bacteria were lysed: while no lytic activity was seen against *K. pneumoniae* at pH 2.5 or higher, for the other four bacteria, a significant level of activity was still observed at pH 2.5 and 3. Only *B. subtilis* was lysed by *D. discoideum* extracts at a pH of 4 or higher.

**Figure 2.**
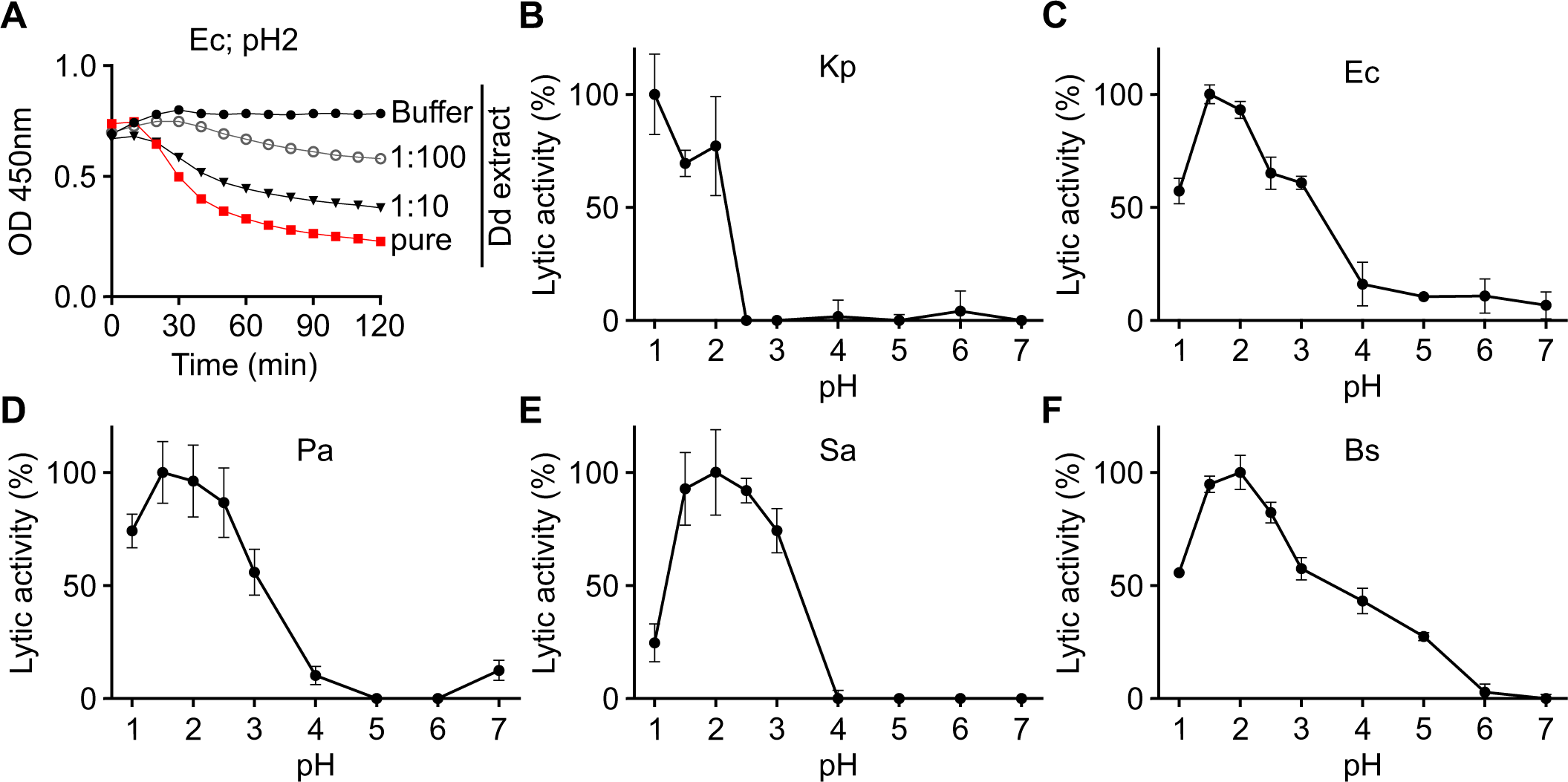
Antibacterial activity of D. discoideum extracts at different pH. **A**. In this example, WT D. discoideum cell extract was serially diluted (from 1/10 to 1/100) in lysis buffer and then mixed with E. coli (Ec) bacteria at pH2. The bacteriolytic activity was monitored over time by spectrophotometry at 450 nm (OD450). **B-F**. Antibacterial activity was determined over a range of pH against K. pneumoniae (Kp) (**B**), E. coli (Ec) (**C**), P. aeruginosa (Pa) (**D**), S. aureus (Sa) (**E**) and B. subtilis (Bs) (**F**). Mean ± SEM; N= 3 independent experiments for Kp, Ec, Sa and Bs; N=5 independent experiments for Pa. The method used to calculate the bacteriolytic activity is detailed in Fig. S5.

### Bacteriolytic activity in extracts from *D. discoideum* mutant cells

We next analyzed the bacteriolytic activity present in extracts from *kil1* KO cells. Kil1 is the main sulfotransferase in *D. discoideum* cells, and *kil1* KO cells show a decreased ability to kill and destroy *K. pneumoniae* bacteria [5, 9], and to a lesser extent, *E. coli* bacteria [4]. As previously reported [6], extracts from *kil1* KO cells exhibited a strongly reduced ability to lyse *K. pneumoniae* bacteria (Fig. 3A). Lysis of *E. coli* and *S. aureus* bacteria by *kil1* KO extracts was also less efficient than with WT extracts, but the lysis was significantly weaker (Fig. 3A). Extracts from *kil1* KO cells lysed *P. aeruginosa* and *B. subtilis* as efficiently as extracts from WT cells (Fig. 3A).

**Figure 3.**
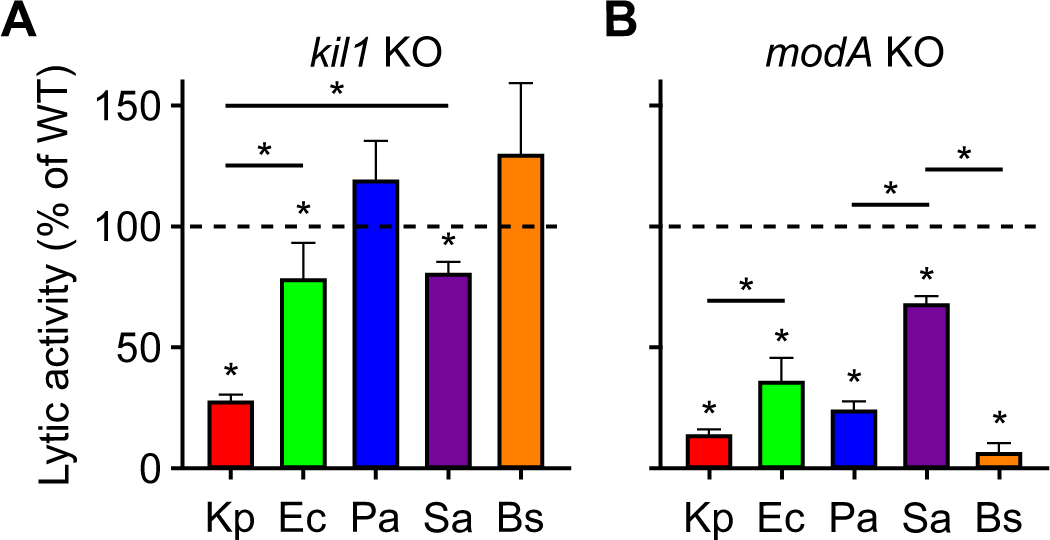
Lytic activity of D. discoideum extracts from kil1 and modA KO cells. The bacteriolytic activity of D. discoideum extracts was tested as described in the legend of Figure 2 at pH 2 against different bacteria. **A**. Extracts from kil1 KO cells exhibited a very reduced activity against K. pneumoniae (Kp) bacteria, a small but significant reduction in activity against E. coli (Ec) and S. aureus (Sa). Antibacterial activity against P. aeruginosa (Pa) and B. subtilis (Bs) was not significantly different in extracts from kil1 KO cells and WT cells. **B**. Extracts from modA KO cells lysed all bacteria less efficiently than extracts from WT cells. The effect was most pronounced against K. pneumoniae, P. aeruginosa and B. subtilis bacteria, and less against E. coli and S. aureus bacteria. Mean± SEM; *: p<0.05; Mann-Whitney test; N=5 independent experiments for Kp and Sa, 9 for Pa and Bs, 10 for Ec exposed to kil1 KO cell extracts; N=5 independent experiment for all 5 bacteria exposed to modA KO cell extracts.

We then tested extracts from a different *D. discoideum* mutant. ModA is a glycosidase present in the ER which plays a key role in the maturation and transport of many lysosomal enzymes. In *modA* KO cells, the activity of many lysosomal enzymes is significantly reduced [10]. We generated *modA* KO cells as detailed in the Material Methods section and in Fig. S1. Extracts from *modA* KO cells lysed all five bacteria species much less efficiently than extracts from WT cells (Fig. 3B). The defect was very strong for lysis of *K. pneumoniae*, *P. aeruginosa* and *B. subtilis*, and less prominent for the lysis of *E. coli* and *S. aureus*.

### Charge-based fractionation of *D. discoideum* bacteriolytic factors

As previously described, a large number of putative lysosomal enzymes can be purified based on their ability to bind to a positively charged resin at a very acidic (pH 3) [6], presumably reflecting the fact that many lysosomal enzymes are decorated with sulfated or phosphorylated sugar moieties, which would remain negatively charged even at a very acidic pH [11]. We used this property to separate *D. discoideum* cell extracts into two main fractions, the fraction bound to anion exchange beads, and the unbound fraction. For the three gram-negative bacteria tested (*K. pneumoniae, E. coli* and *P. aeruginosa*), bacteriolytic activity was detected in the fraction of proteins binding the anion exchange resin and was entirely depleted from the unbound fraction (Fig. 4A-C). On the contrary, bacteriolytic activity against gram-positive bacteria (*S. aureus* and *B. subtilis*) was found in the fraction of proteins not bound to the resin and absent from the fraction of bound proteins (Fig. 4D and E).

**Figure 4.**
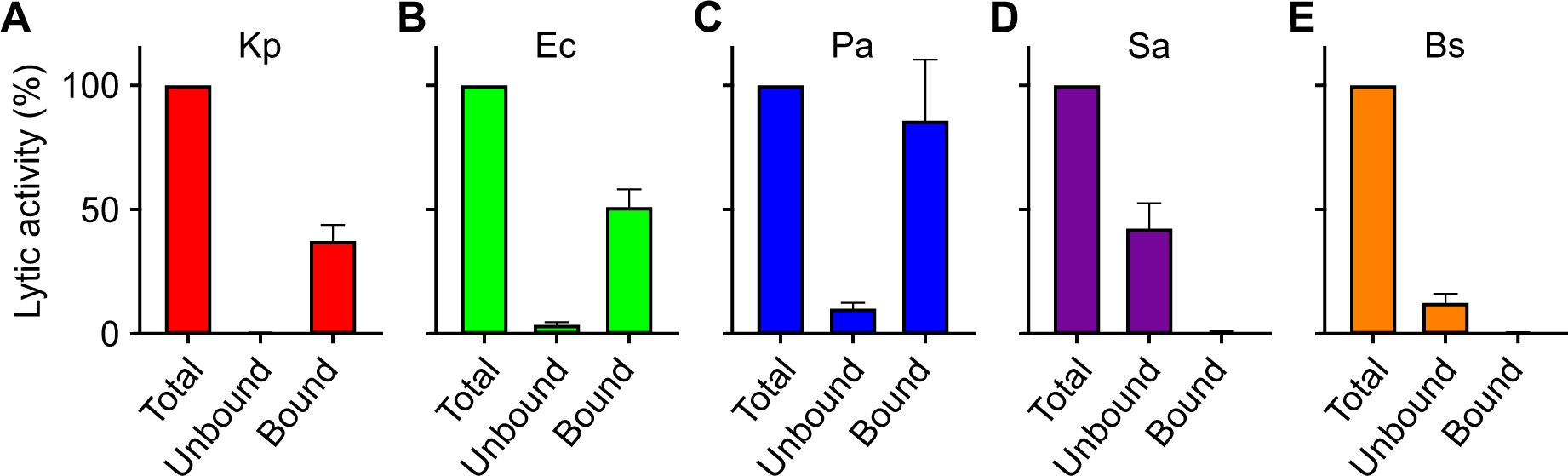
Initial fractionation of D. discoideum extracts. Extracts from D. discoideum (Total) were incubated with an anion exchange resin at pH 3.0. The supernatant (Unbound) was collected. The resin was washed, then the bound proteins (Bound) were eluted in 1M NaCl. The bacteriolytic activity of the different fractions was tested against K. pneumoniae (Kp) (**A**), E. coli (Ec) (**B**), P. aeruginosa (Pa) (**C**), S. aureus (Sa) (**D**) and B. subtilis (Bs) (**E**). Mean ± SEM; N= 6 independent experiments for Kp, Ec, Pa and Sa; N=5 independent experiments for Bs. The antibacterial activity was observed mostly in the Bound fraction for gram-negative bacteria and in the Unbound fraction for gram-positive bacteria.

### Fractionation of bacteriolytic activity against gram-negative bacteria

We then purified further the proteins responsible for the observed bacteriolytic activity against gram-negative bacteria. For this, we eluted the proteins bound to the anion-exchange resin in the presence of increasing concentrations of NaCl and measured the bacteriolytic activity in each eluted fraction (Fig. 5A). The results of six independent experiments were pooled, allowing subtle differences to be reliably detected. Remarkably, the bacteriolytic activities against *K. pneumoniae*, *E. coli* and *P. aeruginosa* were differently distributed in fractions eluted at different salt concentrations (Fig. 5A): the maximal lytic activity was found in the fraction eluted by 200 mM NaCl for *K. pneumoniae*, 250 mM NaCl for *E. coli*, and 300 mM NaCl for *P. aeruginosa*.

**Figure 5.**
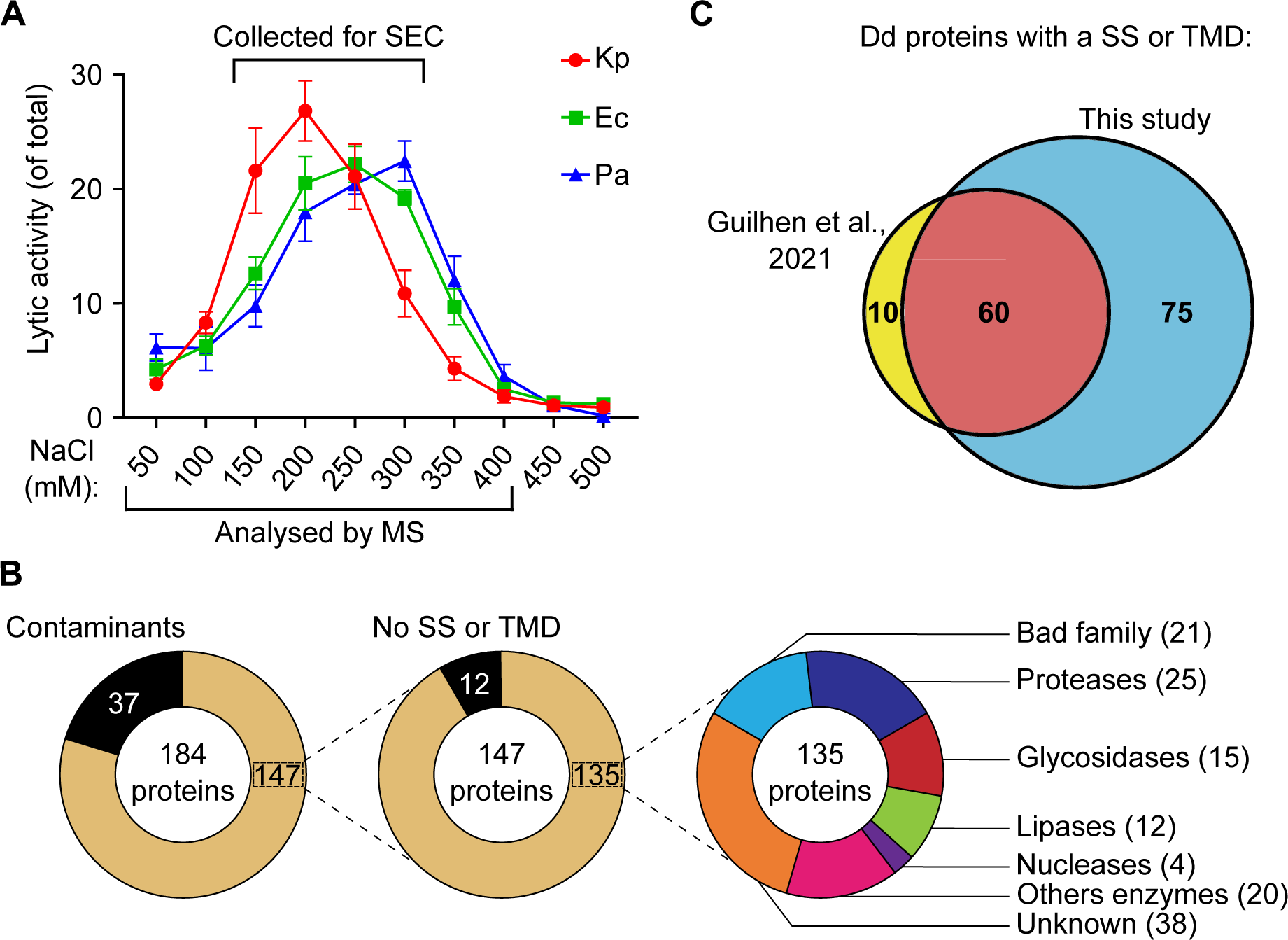
Separation of antibacterial proteins bound to an anion exchange resin. Extracts from D. discoideum were incubated with an anion exchange resin at pH 3. The resin was washed, then the bound proteins were eluted at increasing concentrations of NaCl. (**A**). The antibacterial activity of each fraction was tested against K. pneumoniae (Kp), E. coli (Ec) and P. aeruginosa (Pa). Activity against K. pneumoniae was most prominent in fractions eluted at lower salt concentrations (200 mM NaCl) than activity against E. coli (250 mM) and P. aeruginosa (300 mM). Mean ± SEM; N= 6 independent experiments. (**B**). The protein content of each fraction was analyzed by mass spectrometry and revealed the presence of a large number of presumptive lysosomal enzymes. The detailed list and abundance of different proteins with a signal sequence (SS) or a transmembrane domain (TMD) in each fraction is detailed in Table S1. (**C**). The list of proteins identified in the current study was compared with a similar earlier study [6]. The list of proteins identified in the current study comprised the majority of proteins previously identified (60 proteins in common) plus a large set of previoulsy unidentified proteins (75 proteins). SEC: size exclusion chromatography; Dd: D. discoideum ; MS: mass spectrometry.

We used mass spectrometry to analyze the eight fractions exhibiting bacteriolytic activity (50 mM to 400 mM NaCl). As previously described [6], only a small set of *D. discoideum* proteins attached to an anion exchange resin at acidic pH (Fig. 5B). A total of 147 *D. discoideum* proteins were detected in fractions eluted from the resin. Of these 147 proteins, 135 (92%) exhibited a putative signal sequence (SS) and/or transmembrane domain (TMD). Of these, a large number have a known or suspected lytic activity, and 38 (30%) have no known activity (Fig. 5B). A more detailed analysis of this set of proteins is presented below. This set of proteins includes the vast majority of the proteins detected in a similar previous study [6] (60 proteins), as well as a large number of previously undetected proteins (75 proteins) (Fig. 5C). This is presumably due to the fact that larger amounts of proteins were purified in the current study. A list of the 20 most abundant proteins bound to an anion exchange resin is presented in Fig. S2.

We further purified *D. discoideum* proteins by separating them based on their molecular weight on a gel filtration column. For this we grouped the four fractions eluted at 150-300 mM NaCl and applied them on a Superdex 200 column. The lytic activity against *K. pneumoniae*, *E. coli* and *P. aeruginosa* was then measured in each fraction (Fig. 6A). We observed again that the lytic activities against the three gram-negative bacteria did not co-purify: the lytic activity against *P. aeruginosa* was more abundant in early fractions containing larger proteins (14 mL elution volume, estimated size 70 kDa), while for *E. coli* it peaked in later fractions (15 mL elution volume, ≈ 40 kDa) and for *K. pneumoniae* it was found in even later fractions (15.5-16 mL; ≈ 30 kDa) (Fig. 6A). Experimental variability prevented reliable averaging of multiple experiments, but other experiments showed very similar results (another example is shown in Fig. S3). Proteomic analysis of nine fractions (from 13 to 17 mL) detected the presence of a total of 98 *D. discoideum* proteins with a SS or a TMD. At least 66 of these 98 proteins (67%) exhibited a known or suspected lytic activity (Fig. 6B). This set of proteins is much larger than in a previous study [6] (Fig. 6C). A list of the 20 most abundant proteins purified after size exclusion chromatography is presented in Fig. S4. A selected set of the most abundant proteins identified in this study is shown, including proteases, glycosidases, lipases and membrane-permeabilizing proteins (Fig. 7). Given the fact that bacteriolytic activity was observed in all fractions analyzed, some of these proteins, separately or in combination must be able to lyse gram-negative bacteria. The complete list of all *D. discoideum* proteins with SS or a TMD detected in this study is shown in Table S1.

**Figure 6.**
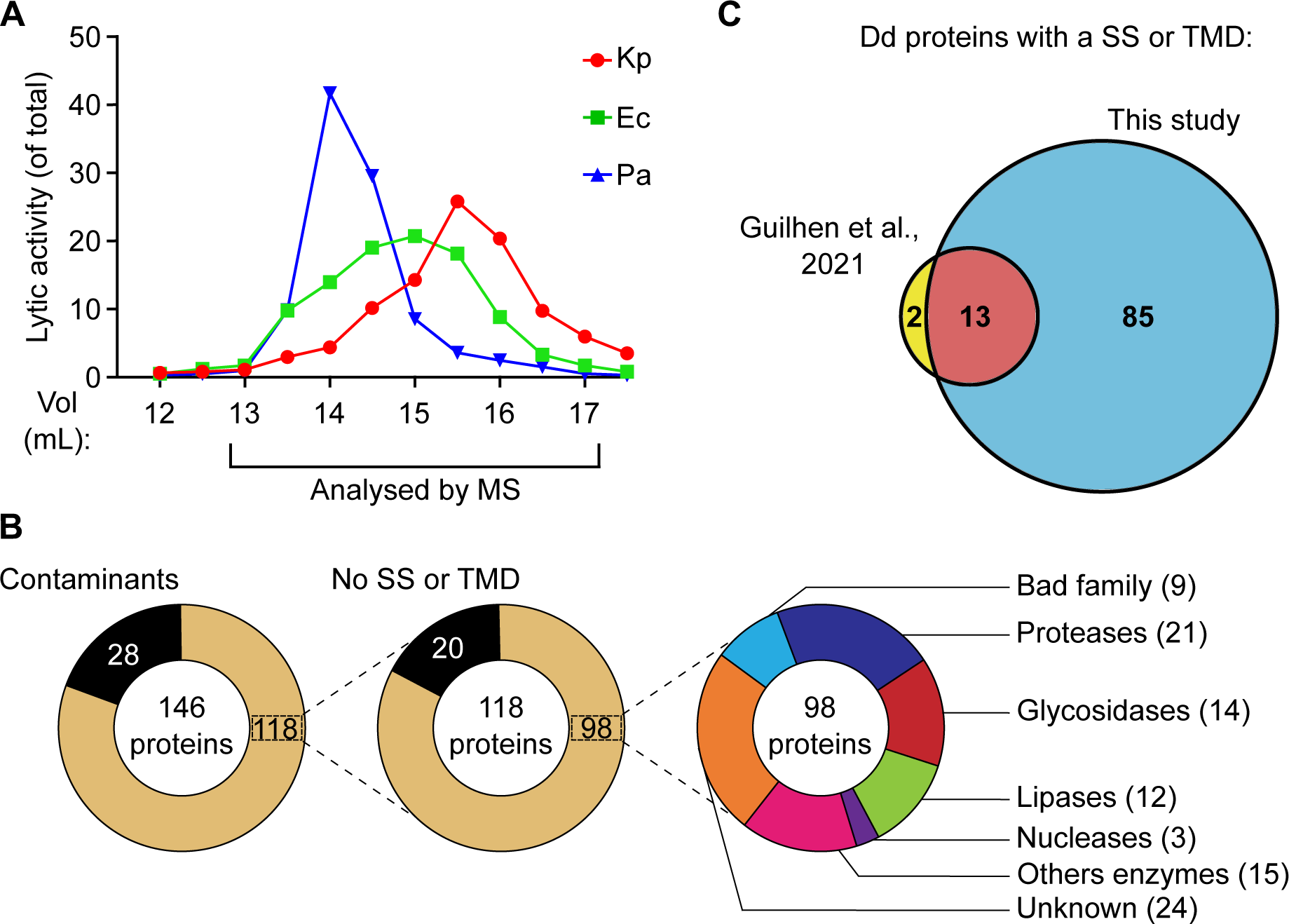
Size exclusion chromatography of antibacterial extracts. Fractions indicated in Fig. 5A were collected, mixed, and further separated on a size exclusion column. (**A**). A representative experiment shows that antibacterial activity against K. pneumoniae (Kp), E. coli (Ec) and P. aeruginosa (Pa) was most prominent in different fractions. Another example is shown in Figure S3. (**B**). The protein content of each fraction was analyzed by mass spectrometry and revealed the presence of a large number of presumptive lysosomal enzymes. The detailed list and abundance of different proteins with a signal sequence (SS) or a transmembrane domain (TMD) in each fraction is detailed in Table S1. (**C**). The list of proteins idenfied in the current study comprised the majority of proteins previously identified (13 proteins in common) [6], plus a large set of previoulsy unidentified proteins (85 proteins). Dd: D. discoideum ; MS: mass spectrometry.

**Figure 7.**
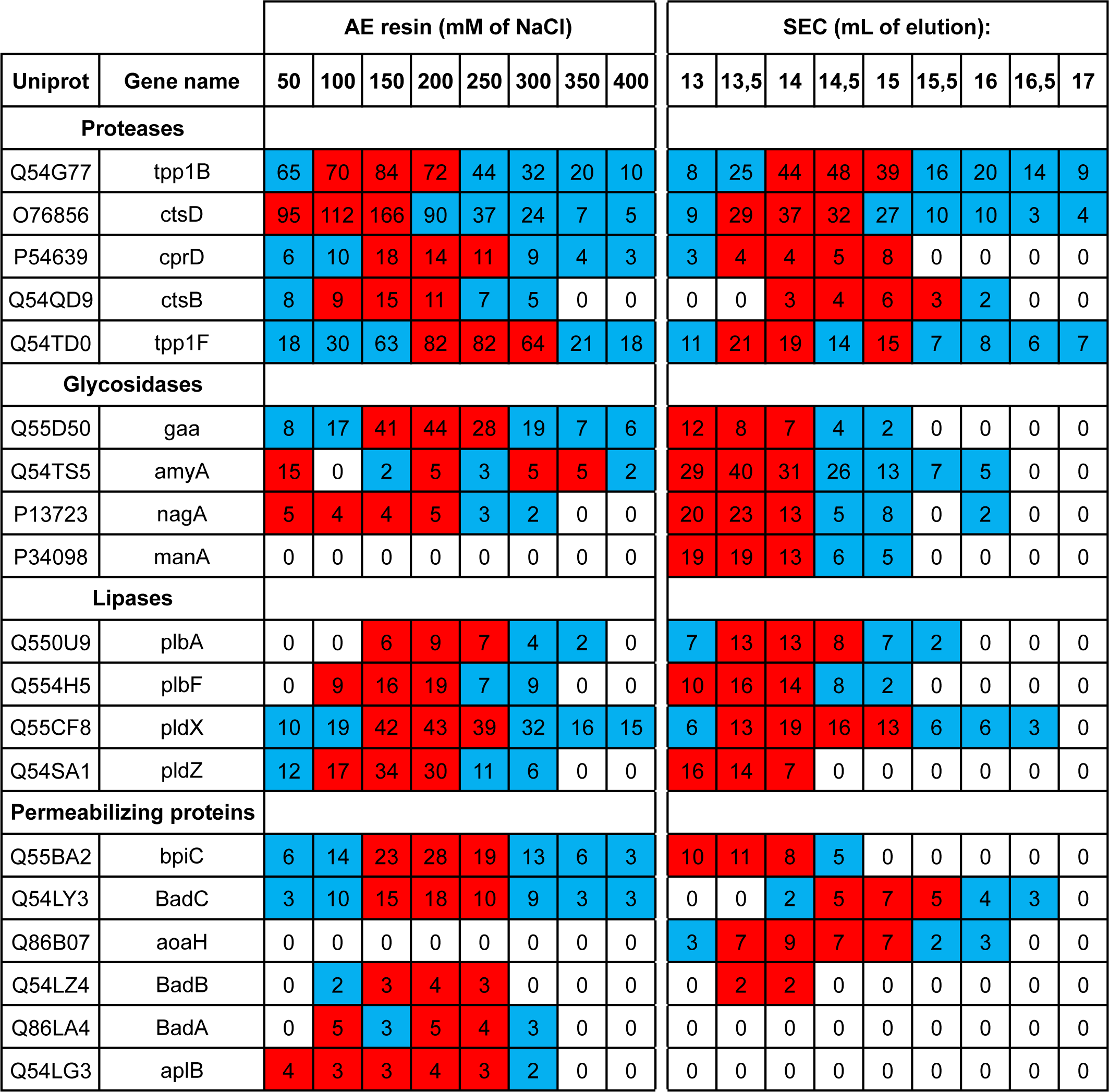
A list of candidate antibacterial effectors. A few putative proteases, glycosidases, lipases and permeabilizing proteins were selected among the most abundant proteins detected in proteomics analysis. For each protein, the Uniprot number, as well as its gene name, and the number of peptides detected by mass spectrometry (spectrum count) in each fraction are indicated. For each protein, the 3 highest spectrum count values are highlighted in red while the other positive spectrum count are in blue. A complete set of all proteins with a signal sequence (SS) or a transmembrane domain (TMD) detected in this study is shown in Table S1. AE: anion exchange; SEC: size exclusion chromatography

## DISCUSSION

In this study, we show that *D. discoideum* extracts exhibits lytic activity against five different bacteria, three gram-negative (*K. pneumoniae, E. coli, P. aeruginosa*) and two gram-positive (*S. aureus* and *B. subtilis*). Together, our results suggest that different molecular mechanisms lead to the lysis of these five bacteria, as summarized graphically in Fig. 8. First, the lytic activity against each bacteria presented a varying degree of susceptibility to pH. The lysis of all five bacteria was optimal at very acidic pH levels. Lysis of *K. pneumoniae* was only observed at a pH of 2 or lower. In contrast, other bacteria were still lysed at higher pH levels: *E. coli*, *P. aeruginosa* and *S. aureus* were lysed at pH 2.5-3, and *B. subtilis* at pH 4 or higher. Second, genetic inactivation of *kil1* strongly reduced the lytic activity against *K. pneumoniae*, decreased moderately the lytic activity against *E. coli* and *S. aureus*, and did not significantly affect the lytic activity against *P. aeruginosa* and *B. subtilis*. Third, genetic inactivation of *modA* produced an entirely different pattern: the remaining activity against *K. pneumoniae*, *B. subtilis* and *P. aeruginosa* bacteria was very low, while it was higher against *E. coli* and *S. aureus*. This indicates varying degrees of dependency to either *kil1* or *modA*-related mechanisms in between all five bacteria. Fourth, the lytic activity against gram-negative bacteria (*K. pneumoniae*, *E. coli* and *P. aeruginosa*) was readily detected in a fraction of proteins bound to an anion-exchange resin, while the activity against gram-positive bacteria (*S. aureus* and *B. subtilis*) was not. Fifth, proteins bound to anion-exchange resin were eluted stepwise, and the optimal bacteriolytic activity against *K. pneumoniae*, *E. coli* and *P. aeruginosa* was found in different fractions. Sixth, proteins were separated further on a gel filtration column, and again optimal activity against *K. pneumoniae*, *E. coli* and *P. aeruginosa* was found in different fractions. Together these results clearly indicate that different proteins are required for optimal lysis of the five bacteria tested here.

**Figure 8.**
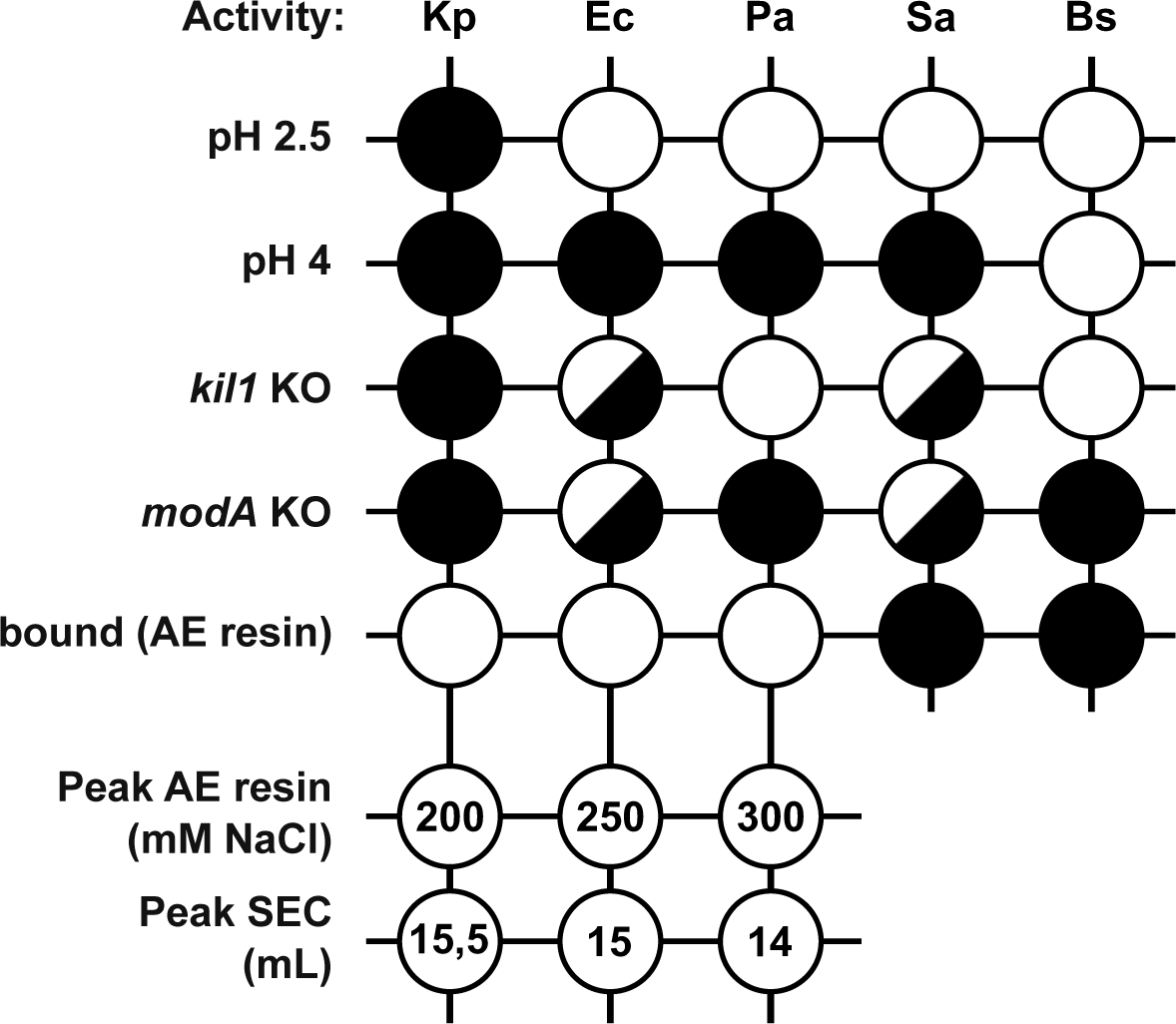
Summary of experimental results. D. discoideum cell extracts lyse all five bacteria tested in optimal conditions (pH 1-2). Differences appeared in other conditions (pH 2.5, pH 4), when extracts from mutant cells (kil1 KO and modA KO) were used, when only proteins binding an anion exchange [21] resin were tested, or when these proteins were further purified by gradual elution from anionic exchange (AE) resin and by size exclusion chromatography (SEC). Overall these results reveal that lysis of each bacteria tested is affected differentially in different conditions, indicating that different effectors are mobilized for optimal lysis of each bacteria. White circle: efficient bacterial lysis; split circle: reduced bacterial lysis; black circle: virtually abolished bacterial lysis. For elution from AE resin and SEC, the fraction containing the highest level of bacteriolytic activity is indicated.

Importantly, in the current study, we did not reach the degree of purity that would allow to assess the bactericidal activity of single proteins, and our attempts to purify further bacteriolytic proteins actually led to a loss of the activity. It is likely that in our experiments bactericidal activity against different bacteria was achieved by the combined activity of several proteins. Our results indicate that different mixtures of bacteriolytic effectors are required to ensure optimal lysis of each bacteria. This interpretation is compatible with results obtained in living cells. Indeed, in living cells, genetic inactivation of genes encoding putative lytic effectors like *bpiC* [4] did reduce and delay bacterial destruction in phagosomes. However, in all instances studied so far, the effect was partial, and the bacteria were eventually destroyed, albeit more slowly. This observation suggests that many proteins are taking part in destruction of bacteria in phagosomes, that none of them is essential and that their effect is cumulative. Although the evidence presented in the current study is very different from previously published evidence, it leads to a very a conclusion. Ultimately, a combination of biochemical and genetic approaches will probably be ideal to fully define the mechanisms at play during bacterial destruction in phagosomes. One of the advantages of biochemical analysis is to identify new proteins potentially implicated in the bacterial destruction in phagosomes. The current study notably provides a short list of proteins potentially implicated in this process, which represent promising candidates for further analysis.

## METHODS

### *D. discoideum* culture and strains

*D. discoideum* DH1-10 parental strain [12] referred to for simplicity as wild-type (WT), as well as *kil1* [9] and *modA* KO (this study) strains were cultured in HL5 medium [13] at 21°C in suspension and diluted twice a week. Cells were collected at a cellular density of 2 to 5 x 10^6^ cells/mL. The *modA* KO cells were created using CRISPR/Cas9 technology as described previously [14]. Briefly, two sgRNA sequences were chosen using the website http://www.rgenome.net/cas-designer/. Each sgRNA was cloned into the pTM1285 plasmid using the Golden-gate assembly and the BpiI enzyme before transformation in TOP10 *E. coli*. Plasmids were purified using a maxiprep kit (Macherey-Nagel, #740574.25) and sequenced. *D. discoideum* cells (16 x 10^6^ cells in 400 μL) were electroporated with the pTM1285/sgRNA plasmid (20 μg) using a Bio-Rad Electroporator (Gene Pulser Xcell™ System)_[15]. Transfected *D. discoideum* cells were resuspended in 35 mL of HL5 medium. The next day, 15 μg/mL of G418 was added to the medium to select plasmid-containing cells. Six days later, surviving *D. discoideum* cells were cloned by limiting dilution in the absence of G418 in 96-well plates. To analyze individual clones, the genomic region of interest was amplified by PCR and sequenced. Two clones presenting different mutations were kept for further analysis. The detailed description of the *modA* KO mutant generated and analyzed in this study is found in Fig. S1.

### Bacterial culture and strains

The bacterial strains used were *K. pneumoniae* KpGe [16], *E. coli* REL606 [17], *P. aeruginosa* PT531 [18], nonsporulating *B. subtilis* strain 36.1 [19], and *S. aureus* ATCC 29213. Bacteria were grown overnight in LB (lysogeny broth) agar plates at 37°C. Before each experiment, a single colony was picked from the plate, inoculated into 3 mL of liquid culture medium (LB for *K. pneumoniae* and *S. aureus*, SM (standard medium) for *P. aeruginosa* and *B. subtilis* and SM supplemented with 10 g/L of glucose for *E. coli*) [13] and grown for 16 h under agitation at 37°C.

### Bacterial lysis assessed by measuring optical density

To measure bacterial lysis, 1× 10^8^ *D. discoideum* cells were washed twice in phosphate buffer (PB: 2 mM of Na2HPO4 and 14.7 mM of KH2PO4, pH 6.3) and lysed in 1mL of Lysis buffer (50 mM NaPO_4_ buffer, pH5, 0.5% Triton X-100) for 20 min at 4°C with occasional vortexing (20-30 sec, every 5 min). The cell lysate was centrifuged (30,000 g, 20 min at 4°C) to remove unlysed cell debris and nuclei and the supernatant collected. The cleared supernatant was buffered at the desired pH using HCl 1M or NaOH 5N, then diluted using lysis buffer at the desired pH (from 1 to 7). Bacteriolytic activity was assessed by mixing in a 96 well microtiter plate 100 μL of cell extract (or Lysis buffer as a negative control) with 100 μL of bacterial culture washed once in 50 mM NaP buffer at the corresponding pH and resuspended to a final optical density at 450 nm of 0.8-1.2. The decrease in turbidity at 21°C (optical density at 450 nm) was measured every 10 min for 2 h, with a spectrophotometer plate reader (Hidex Sense Plate Reader, Software 1.3.0). To calculate the bacteriolytic activity of a sample, the OD_450_ values were normalized to the value of the sample at time 0 defined as 100%. The normalized OD_450_ values were then plotted as a function of time and the area under the curve (AUC) was calculated. The activity was then calculated by comparing with a standard curve obtained by diluting WT *D. dicoideum* cell lysates. The calculation method used is detailed in Fig. S5.

### Bacterial lysis assessed by direct observation

To visualize bacterial lysis, bacteria were exposed to *D. discoideum* cell extracts as described above for 2 h, then 4 µL were placed on a glass slide and covered with a glass coverslip. The bacteria were imaged using a Zeiss Axio Observer Z1 with Definite Focus 2 microscope and a EC Plan-Neofluar 20x / 0.50, Ph2 WD 2.0 mm objective. To quantify the degree of lysis, raw TIFF phase contrast images were processed using MATLAB R2023a (The MathWorks). Initially, a flat-field correction was applied, followed by a non-local mean filter on all images. Subsequently, these processed images were analyzed using QuPath 0.5.0 [20] where an artificial neural network was employed to classify pixels corresponding to bacteria within the field of view. This automated detection of bacteria was also corrected manually. For each sample, 3 separate images (each 624 x 501 μm), were analyzed to determine the area covered by bacteria.

### Biochemical fractionation

*D. discoideum* extracts were fractionated essentially as described [6]. Briefly, *D. discoideum* cleared cell lysates described above were buffered at pH 3, then mixed with 200 µL of anion exchange resin (Q Sepharose Fast Flow; Sigma #17-0510-10) pre-washed in Lysis buffer at pH 3. After 1 h of incubation at 4°C on a rotating wheel, the anionic resin was pelleted (30,000 g for 30 sec at 4°C), and the supernatant collected (Unbound fraction). The resin was then washed 5 times with 1 mL of Lysis buffer at pH 3. Negatively charged proteins attached to the resin were eluted at pH 3 in lysis buffer containing 1M of NaCl (Bound fraction). The bacteriolytic activity of total cell extracts, unbound and bound fractions were then assessed as described above.

To purify further the bacteriolytic activity, a more concentrated cell extract was used (5×10^8^ *D. discoideum* cells lysed in 1.75 mL of Lysis buffer) and negatively charged proteins attached to the resin were eluted at increasing concentrations of NaCl, from 50 to 500 mM of NaCl with increments of 50 mM. Recovered fractions were tested for the presence of bacteriolytic activity. Fractions presenting the highest bacteriolytic activity (e.g. fractions 150, 200, 250 and 300 mM of NaCl) were mixed, buffered at pH 7 and loaded on a size-exclusion chromatographic (SEC) column (Superdex 200 10/300 GL; Sigma #17-5175-01). The column was equilibrated with NaP buffer at pH 7 and eluted with the same buffer at a rate of 18 mL/h. Fractions of 0.5 mL were collected and their bacteriolytic activity tested.

### Protein identification by ESI-LC-MSMS

Protein samples (50 μL) were dried, resuspended in Laemmli buffer and purified by 1D gel electrophoresis (SDS-PAGE; stacking gel method) using an home-made 4% bis-acrylamide stacking gel and a 12% bis-acrylamide resolving gel. Briefly, samples were loaded, and migration was performed at 70 V until the proteins enter the resolving gel. ((The gel was stained with SilverQuest silver (Life technologies). Bands of interest were cut, destained with the SilverQuest destainer solutions and In-gel digestion was performed as follow: gel pieces were incubated in 100 μL of 50% acetonitrile (AcN) in 50 mM ammonium bicarbonate (AB) for 15 min at room temperature. Proteins were reduced by incubation of gel pieces for 30 min. at 50°C in 100 μL of 10 mM DTT in 50 mM AB. The DTT solution was then replaced with 100 μL of 55 mM iodoacetamide in 50 mM AB and proteins were alkylated by incubation of the gel pieces for 30 min at 37°C in the dark. Gel pieces were then washed for 15 min with 100 μL of 50mM AB and for 15 min with 100 μL of 100% AcN. Gel pieces were then air dried for 15 min at room temperature. Dried pieces of gel were rehydrated for 45 min at 4°C in 30 μL of a solution of 50 mM AB containing trypsin at 10 ng/ μL and 0.01% of Protease Max (PM) Surfactant trypsin enhancer (Promega). Subsequently, 10 μL of 0.01% of PM was added before incubating the samples for 1 hour at 50°C. The supernatant was transferred to a new polypropylene tube and an additional peptide extraction was performed with 50 μL of 20% FA (formic acid) for 15 min at room temperature with occasional shaking. Extractions were pooled and completely dried under speed vacuum.

Samples were diluted in 12 μL of loading buffer (5% CH3CN, 0.1% FA) and 4 μL were injected on a column. LC-ESI-MS/MS was performed on a Q-Exactive HF Hybrid Quadrupole-Orbitrap Mass Spectrometer (Thermo Scientific) equipped with an Easy nLC 1000 liquid chromatography system (Thermo Scientific). Peptides were trapped on an Acclaim pepmap100, C18, 3 μm, 75 μm x 20 mm nano trap-column (Thermo Scientific) and separated on a 75 μm x 250 mm, C18, 2 μm, 100 Å Easy-Spray column (Thermo Scientific). The analytical separation was run for 90 min using a gradient of H_2_O/FA 99.9%/0.1% (solvent A) and CH_3_CN/FA 99.9%/0.1% (solvent B). The gradient was run as follows: 0-5 min 95 % A and 5 % B, then to 65 % A and 35 % B for 60 min, and 5 % A and 95 % B for 20 min at a flow rate of 250 nL/min. Full scan resolution was set to 60’000 at m/z 200 with an AGC target of 3 x 10^6^ and a maximum injection time of 60 ms. Mass range was set to 400-2000 m/z. For data dependent analysis, up to twenty precursor ions were isolated and fragmented by higher-energy collisional dissociation HCD at 27% NCE. Resolution for MS2 scans was set to 15’000 at m/z 200 with an AGC target of 1 x 105 and a maximum injection time of 60 ms. Isolation width was set at 1.6 m/z. Full MS scans were acquired in profile mode whereas MS2 scans were acquired in centroid mode. Dynamic exclusion was set to 20 s.

Peak lists (MGF file format) were generated from raw data using the MS Convert conversion tool from ProteoWizard. The peaklist files were searched against the Dictyostelium discoideum Reference Database (Uniprot, release 2020-05, 12746 entries) combined with an in-house database of common contaminant using Mascot (Matrix Science, London, UK; version 2.5.1). Trypsin was selected as the enzyme, with one potential missed cleavage. Precursor ion tolerance was set to 10 ppm and fragment ion tolerance to 0.02 Da. Carbamidomethyl of cysteine was specified as fixed modification. Deamidation of asparagine and glutamine, as well as oxidation of methionine were specified as variable modifications. The Mascot search was validated using Scaffold 4.11.1 (Proteome Software). Peptide identifications were accepted if they could be established at greater than 74.0 % probability to achieve an FDR less than 0.1 % by the Scaffold Local FDR algorithm. Protein identifications were accepted if they could be established at greater than 99.0 % probability to achieve an FDR less than 1.0 % and contained at least 2 identified peptides. Protein probabilities were assigned by the Protein Prophet algorithm [21]. Proteins that contained similar peptides and could not be differentiated based on MS/MS analysis alone were grouped to satisfy the principles of parsimony.

## LIST OF ABBREVATIONS

Dd: *D. discoideum*;
Kp: *K. pneumoniae*;
Ec: *E. coli*;
Pa: *P. aeruginosa*;
Sa: *S. aureus*;
Bs: *B. subtilis*;
OD: optical density;
AE: anion exchange;
MS: mass spectrometry;
SEC: size exclusion chromatography;
AUC: area under the curve;
SS: signal sequence;
TMD: transmembrane domain

## DECLARATIONS

### Ethics approval and consent to participate

Not applicable.

### Consent for publication

Not applicable.

### Availability of data and material

The datasets used and/or analyzed during the current study are publicly available on an open access server (Yareta) using this DOI: 10.26037/yareta:oro3nohevfcglcmhyd3hoiefkm .

### Author’s contributions

OL, RM and PC conceived and designed the study. OL, RM, TJ, AB, and CG performed experiments. OL and RM analyzed experiments. OL, RM and PC wrote the manuscript. All authors read and contributed to the manuscript.

---

**Figure S1.**
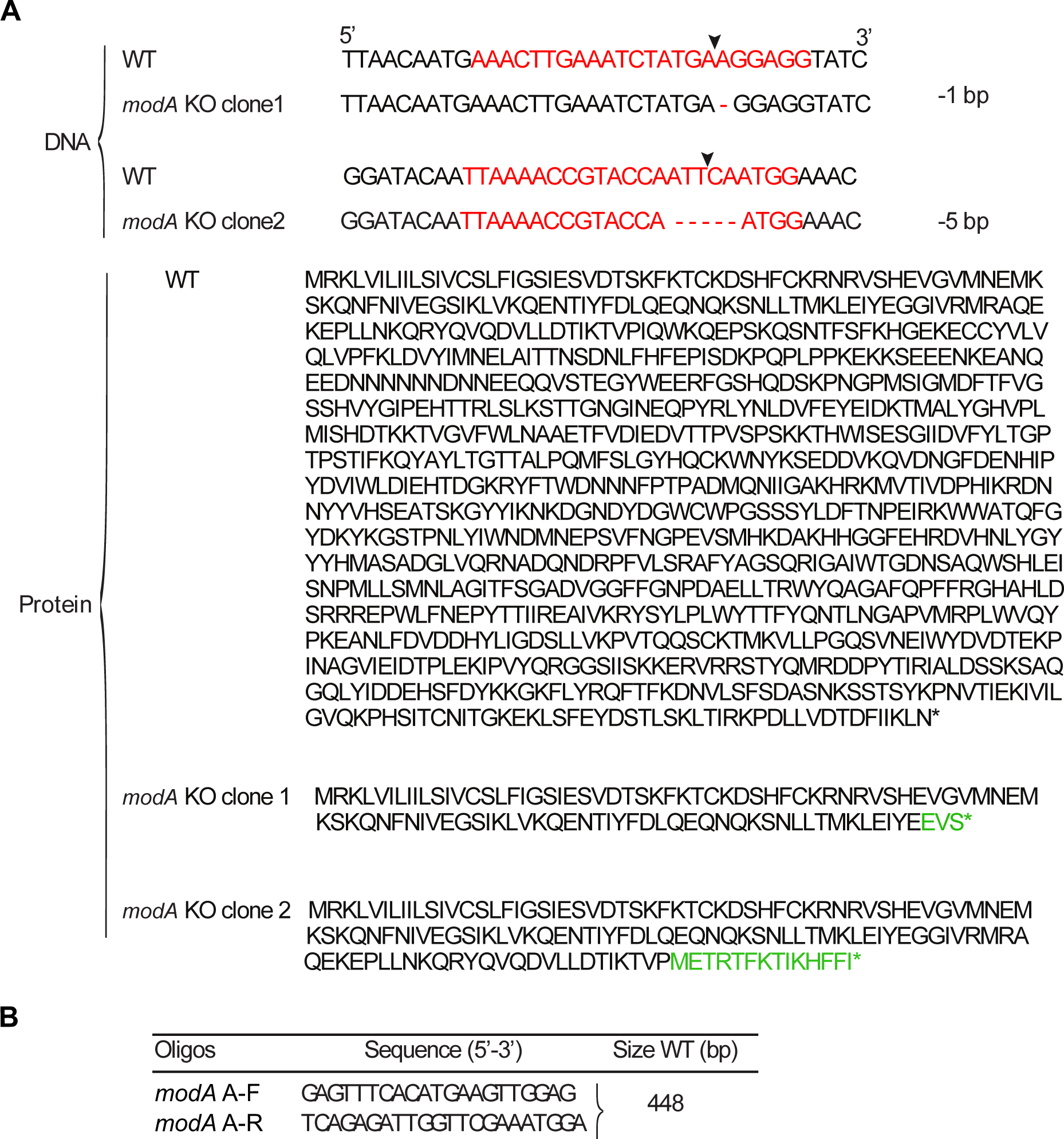
Generation of modA KO cells using CRISPR/Cas9. (**A**) Schematic representation of the modA gene and protein in WT and in KO cells. The sequence targeted by the guide RNA is indicated in red and the arrow indicates the cutting site of the Cas9 nuclease. The sequences of two independent guide RNAs and mutant clones are represented. The truncated and aberrant proteins predicted to be produced in mutant clones are showon. (**B**) Oligonucleotides used for the PCR amplification of the targeted genomic region.

**Figure S2.**
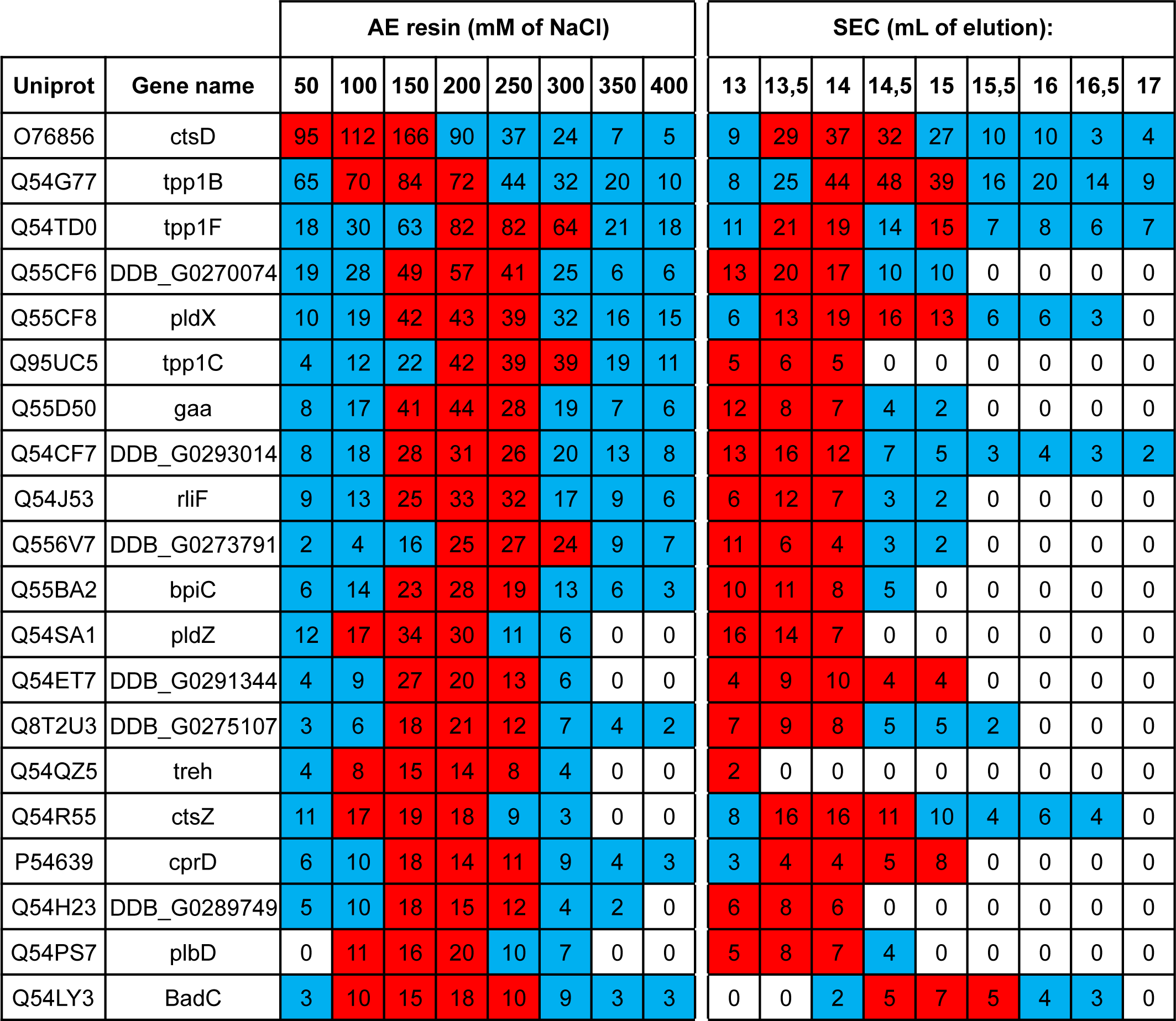
Most abundant proteins bound to an anion exchange resin. The 20 most abundant D. discoideum proteins with a signal sequence (SS) or a transmembrane domain (TMD) binding an anion exchange resin (Fig. 5) are listed. For each protein, the Uniprot number, as well as its gene name, and the number of peptides detected by mass spectrometry (spectrum count) in each fraction are indicated. For each protein, the 3 highest spectrum count values are highlighted in red while the other positive spectrum count are in blue. AE: anion exchange; SEC: size exclusion chromatography

**Figure S3.**
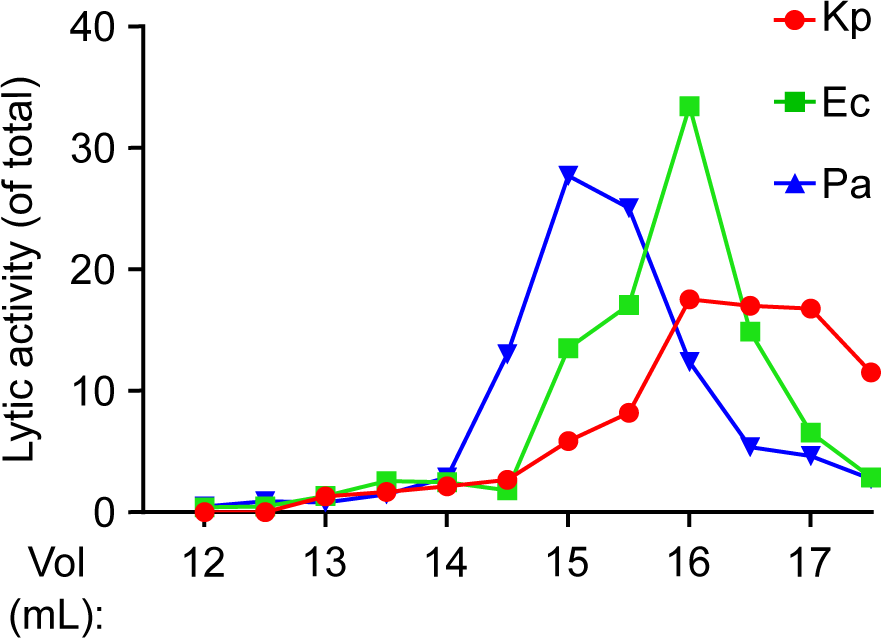
Size exclusion chromatography of D. discoideum extracts. This figure presents an experiment identical to the experiment shown in Fig. 6A. Like the data presented in Fig. 6A, this experiment indicates that the peak of antibacterial activity against K. pneumoniae (Kp), E. coli (Ec) and P. aeruginosa (Pa) was observed in different fractions.

**Figure S4.**
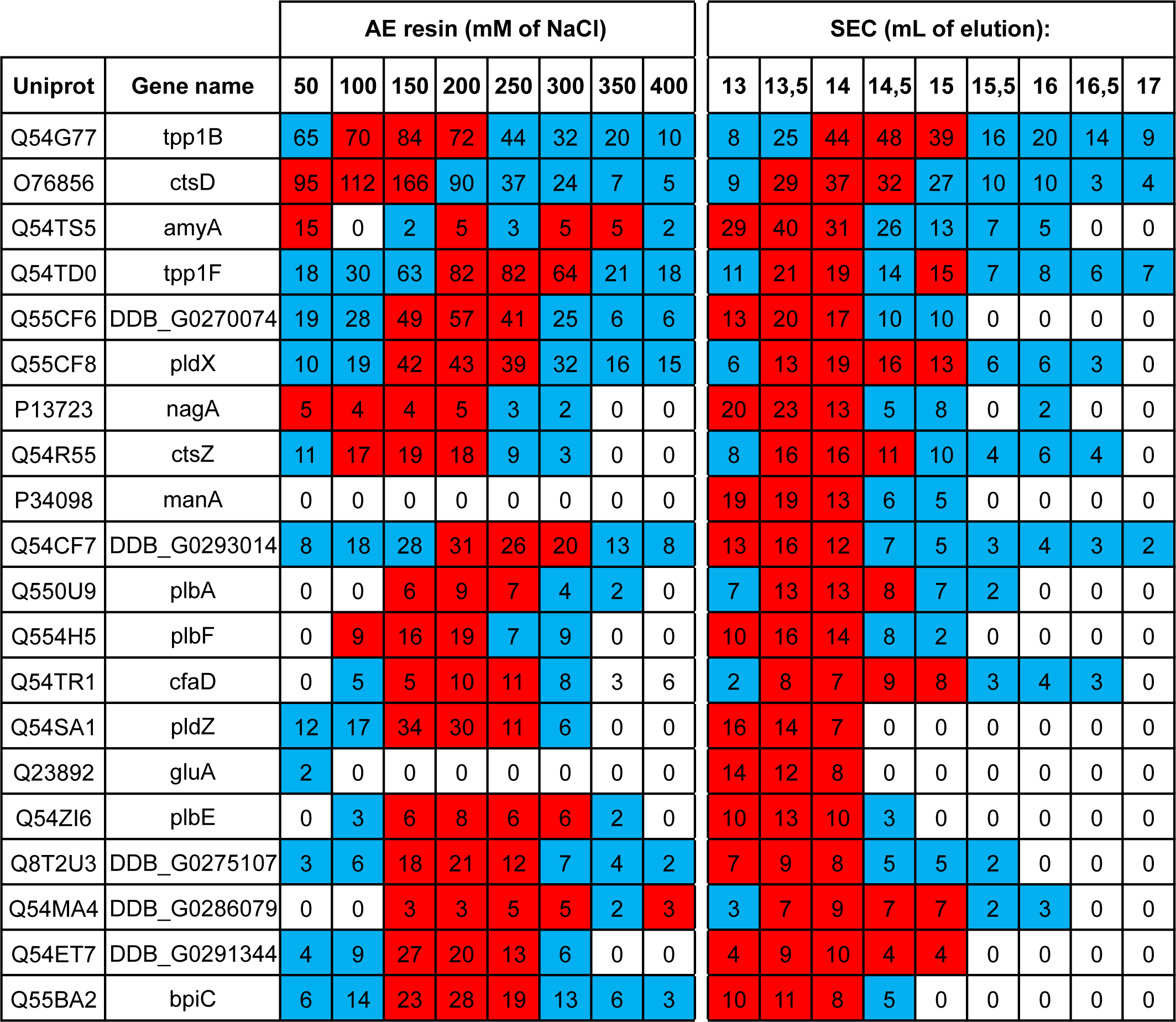
Most abundant proteins collected after separation on a size exclusion column. The 20 most abundant proteins after purification by size exclusion chromatography (Fig. 6) are listed. For each protein, the Uniprot number, as well as its gene name, and the number of peptides detected by mass spectrometry (spectrum count) in each fraction are indicated. For each protein, the 3 highest spectrum count values are highlighted in red while the other positive spectrum count are in blue. AE: anion exchange; SEC: size exclusion chromatography

**Figure S5.**
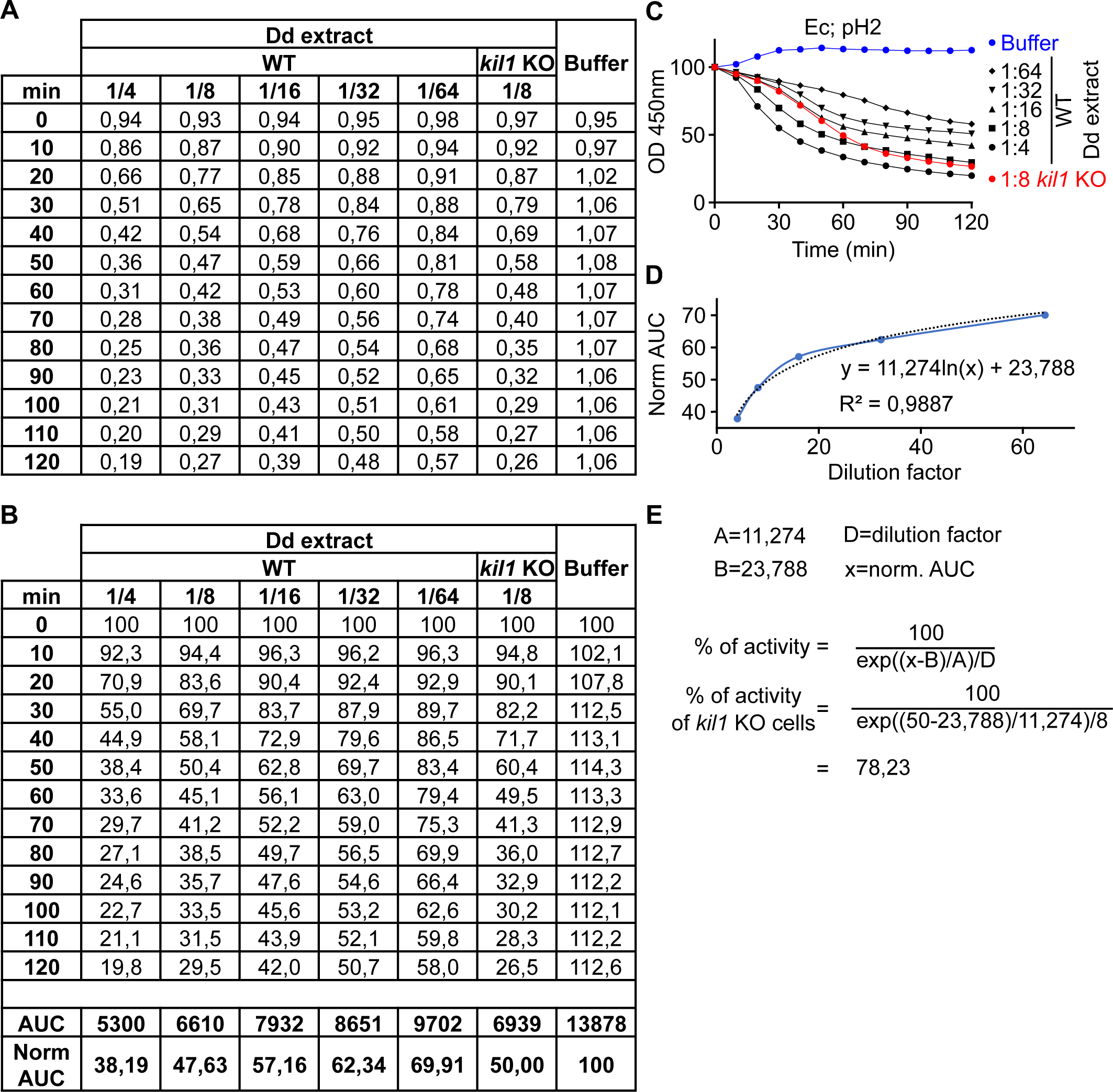
Determination of bacterial lysis activity. (**A**) Raw optical density (OD450nm) values were measured every 10 min over 2 h for a sample of interest (here kil1 KO cell lysate), a standard range of dilution of WT cell lysate and a control condition (buffer). Raw OD450nm values were normalized by setting the value of each sample at time 0 to 100% (**B**) and plotted into curves (**C**). Area under the curves (AUC) were calculated and normalized (“Norm AUC”) by dividing them by the AUC of the buffer condition. (**D**) Norm AUCs of the standard WT cell lysate samples were plotted to generate a calibration curve. (**E**) The equation fitting this standard curve was determined and used to calculate the percentage of activity of the sample of interest (here 78%).

**Table S1.**
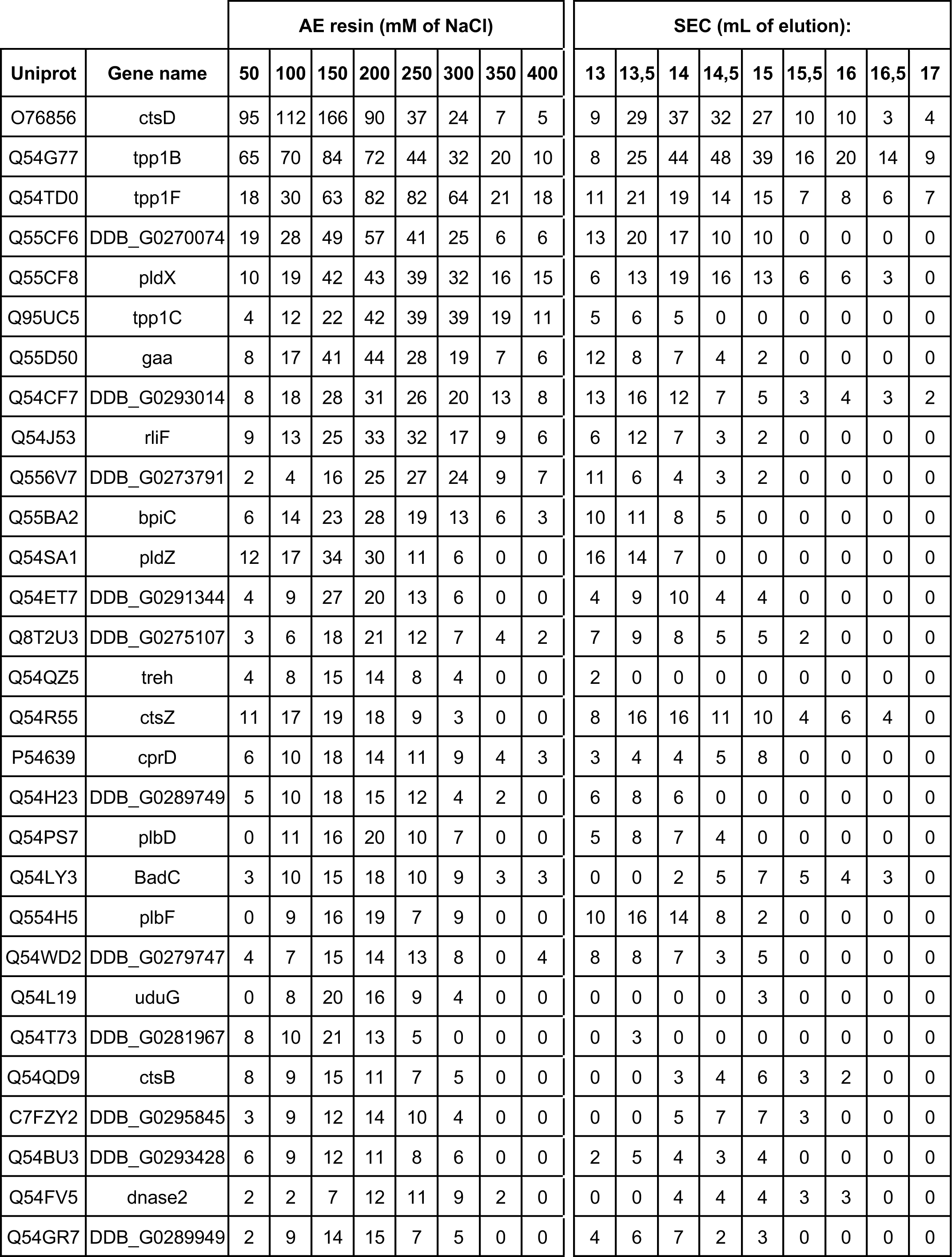
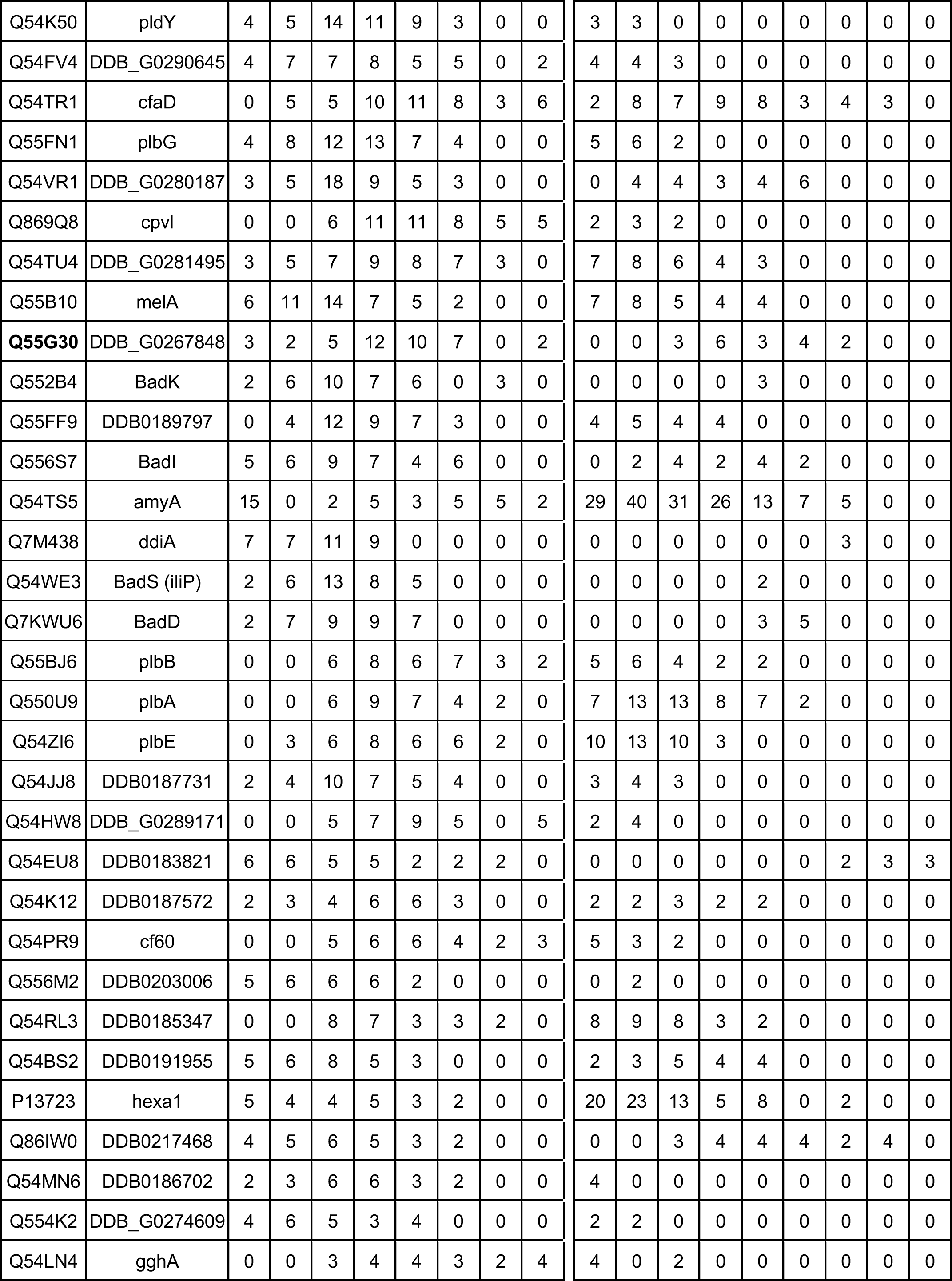
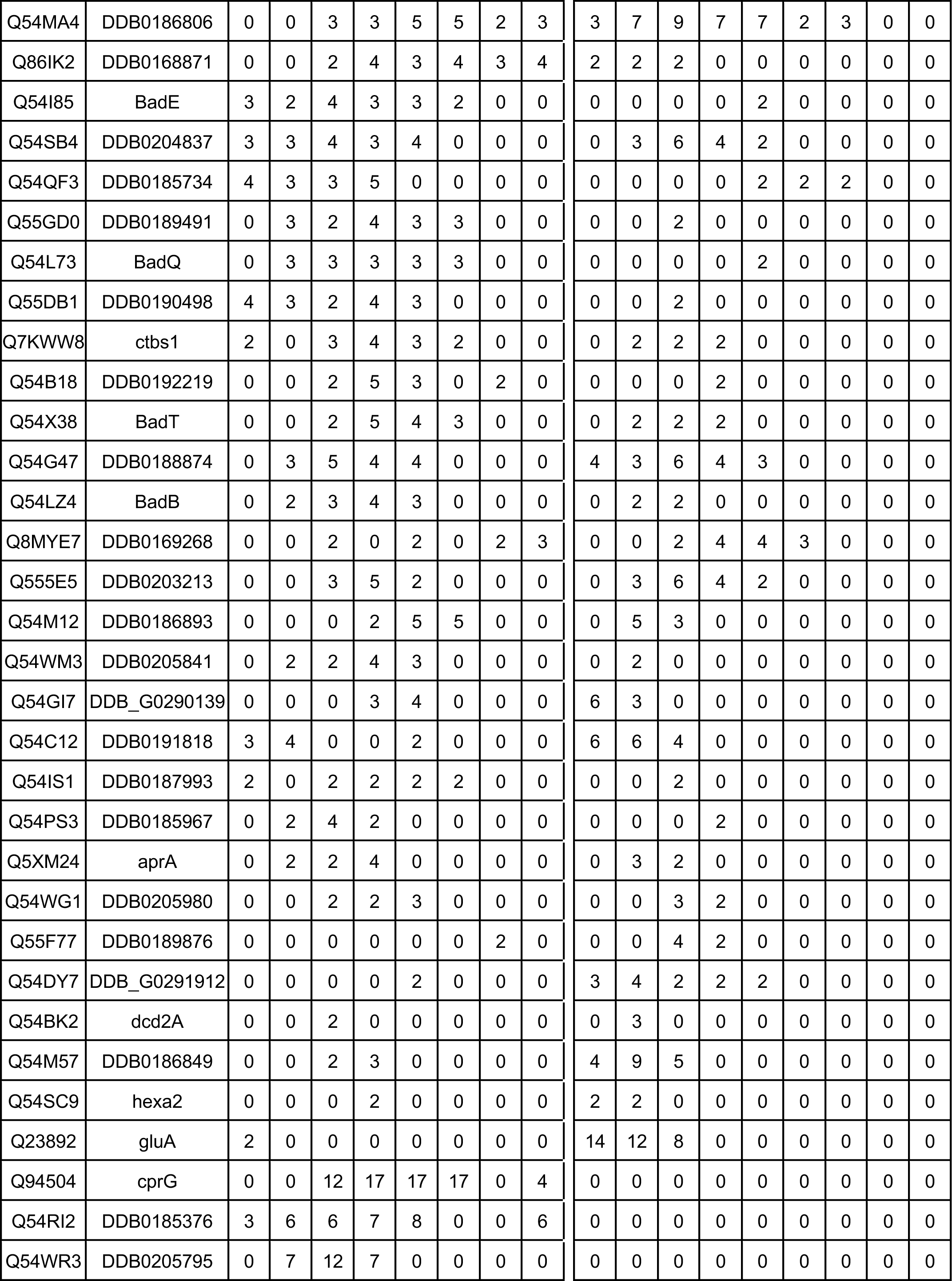
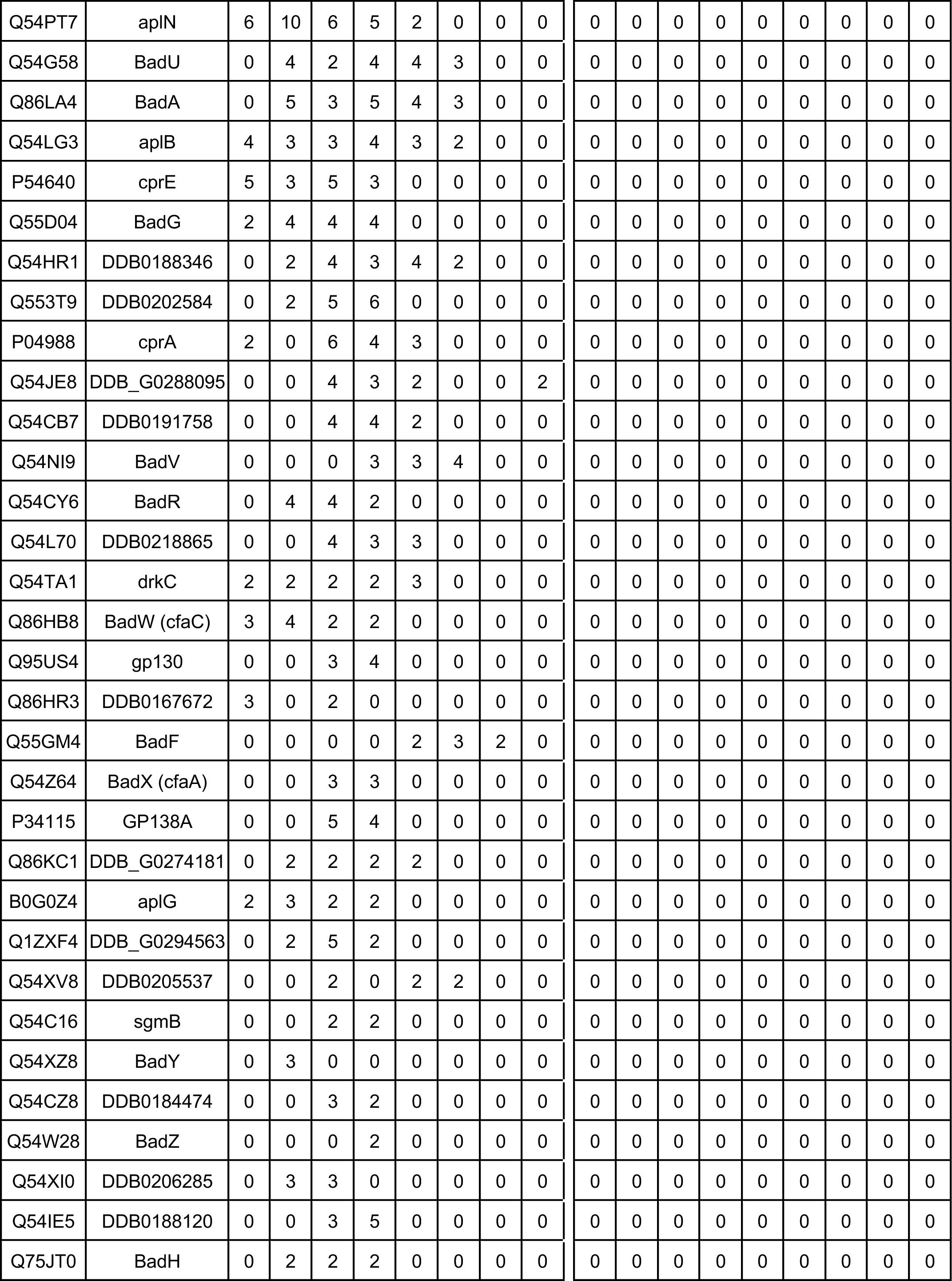
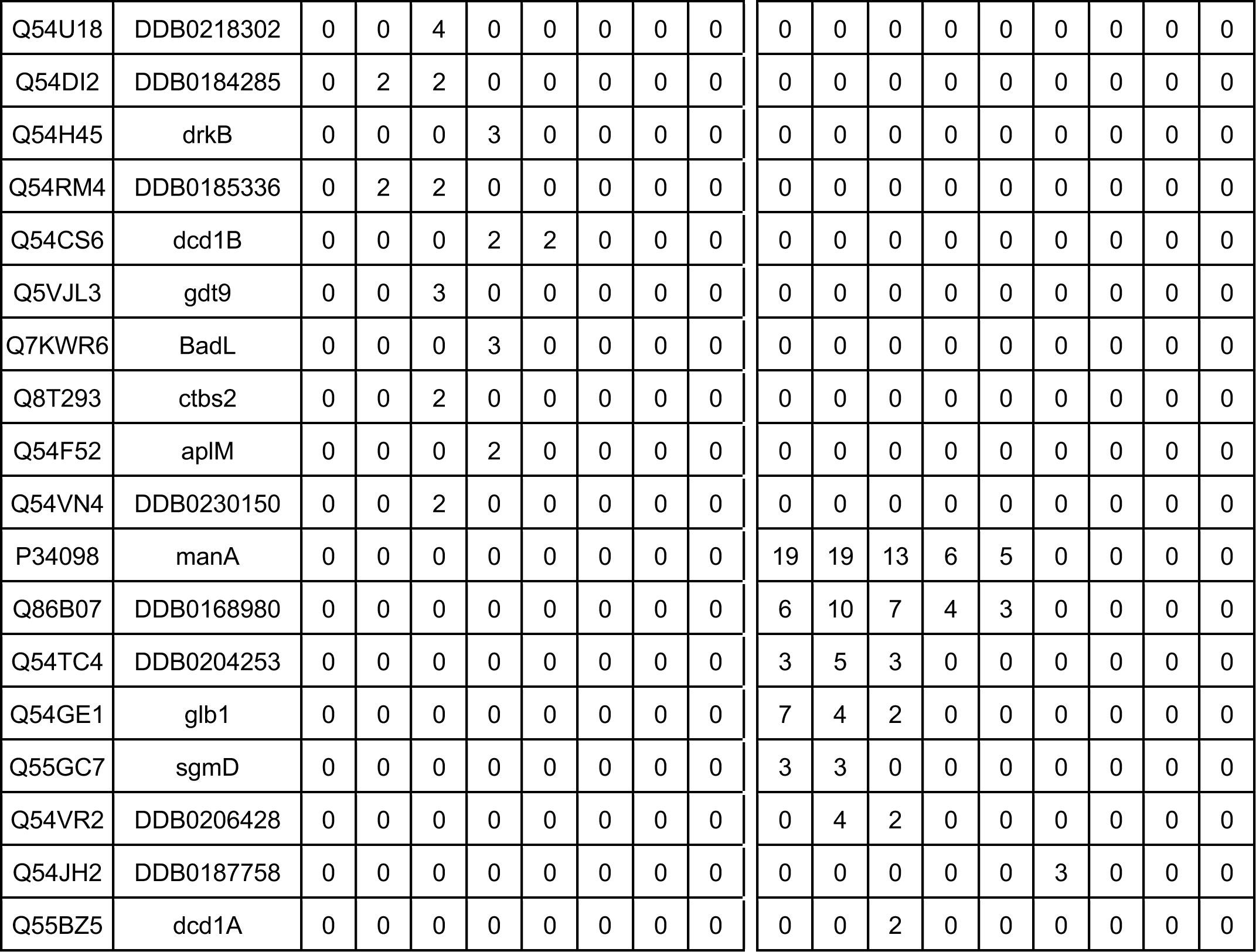
Full list of proteins with a SS or TMD detected in this study. A list of all proteins detected with a signal sequence (SS) or transmembrane domain (TMD) in this study is provided. For each protein, the Uniprot number, as well as its gene name, and the number of peptides detected for a given protein (spectrum count) by mass spectrometry in a fraction are indicated.s

